# Nucleoporin 50a interacts with geminivirus C4 proteins and contributes to infection

**DOI:** 10.64898/2026.08.16.745072

**Authors:** Aparna Chodon, Pandi Gopal, Rosa Lozano-Durán

## Abstract

Geminiviruses are plant DNA viruses that replicate in the nucleus of the host cell and rely on the host nucleocytoplasmic transport machinery to complete their infection cycle. While various geminiviral proteins have been reported to interact with plant transport factors, the contribution of nuclear pore complex components to geminivirus infection remains largely unexplored. Here, we identify nucleoporin 50a (NUP50a) as a previously unreported host factor that contributes to bhendi yellow vein mosaic virus (BYVMV) infection. Affinity purification coupled with mass spectrometry isolated NUP50a as a potential interactor of the BYVMV pathogenicity determinant C4, which was further validated by pull-down and co-immunoprecipitation assays. Yeast two-hybrid assays, bimolecular fluorescence complementation, and colocalization analysis demonstrated that BYVMV C4 directly associates with NUP50a predominantly in the nucleus. Virus-induced gene silencing of *NbNUP50a* significantly delayed symptom development and reduced viral DNA accumulation, suggesting that NUP50a is required for efficient BYVMV infection. Silencing *NbNUP50a* did not influence the subcellular localization of BYVMV C4, indicating that the role of NUP50a extends beyond determining C4 steady-state localization. Notably, NUP50a was found to associate with C4 proteins from three additional geminiviruses, supporting the possibility that targeting NUP50a represents a characteristic strategy among geminiviruses. Together, our findings provide evidence of a nuclear pore complex member involved in geminivirus pathogenesis. These results establish a framework for further study of the potential transport-dependent and/or transport-independent functions of NUP50a during viral infection.

## 1. INTRODUCTION

Viruses are obligate intracellular pathogens that strictly depend on the host cellular machinery to complete their infection cycle. Successful viral infection requires coordinated manipulation of multiple host pathways, which frequently include nuclear transport pathways that mediate the movement of viral and host factors between cytoplasm and nucleus. These nucleocytoplasmic trafficking events are primarily regulated by the nuclear pore complex (NPC), the major gateway controlling the exchange of macromolecules between these cellular compartments. The NPC is one of the largest protein complexes in eukaryotic cells, forming ring-shaped channels that perforate the nuclear envelope (1). In addition to mediating nucleocytoplasmic transport, several NPC components have emerged as important regulators of host antiviral responses. For instance, in *Drosophila*, the virus-induced nucleoporin (NUP) 98 relocalizes from the NPC to the nucleoplasm, where it binds the promoters of antiviral genes and contributes to defense against infection by several human arboviruses (2).

Nucleocytoplasmic shuttling is particularly relevant to geminiviruses, a family of circular single-stranded DNA plant viruses that infect many agroeconomically important crops. Since geminiviruses replicate within the nucleus of infected cells, they depend heavily on host nuclear transport pathways to mediate the import and export of viral components across the nuclear envelope (3). Consistent with this, several geminiviral proteins have been shown to interact with host nuclear transport factors. For example, the coat proteins of tomato yellow leaf curl virus (TYLCV) (*Begomovirus coheni*) and mungbean yellow mosaic virus (MYMV) (*Begomovirus vignaradiatae*) interact with karyopherin-α1 and importin-α, respectively, to facilitate nuclear import (4,5). Similarly, exportin-α mediates nuclear export of the tomato leaf curl Yunnan virus (TLCYnV) (*Begomovirus solanumflavusyunnanense*) C4 protein, thereby promoting its function as a pathogenicity and symptom determinant at the plasma membrane (6). Despite the central role of nucleocytoplasmic trafficking during geminivirus infection, direct roles of nuclear pore complex components in geminivirus infection remain largely unexplored.

NUP50 (Nup2 in yeast) is a dynamic FG-repeat nucleoporin associated with the nuclear basket of the NPC, whose FG-repeat domains function as selective transport barriers within the NPC (1). In animals and yeast, NUP50 regulates nuclear import through interactions with importins and Ran GTPases (7–9). In *Arabidopsis*, NUP50a interacts with NUP50b and several nucleocytoplasmic transport factors, including importins and Ran proteins, supporting a conserved role in nuclear transport (10). AtNUP50a and AtNUP50b were also shown to regulate seed development through transcriptional control of cell wall-associated genes, with loss-of-function mutants exhibiting reduced seed longevity and salinity stress tolerance (11). Furthermore, although NUP50a was initially described as a predominantly nucleoplasmic protein (10), recent studies in *Physcomitrium patens* and other land plants identified its localization at plasmodesmata, as part of a potential plasmodesmata pore complex (PDPC) (12,13). Together, these studies suggest that NUP50a may perform diverse functions beyond its conventional role in nucleocytoplasmic transport, raising the possibility that it could influence host-virus interactions in different ways.

Among the canonical geminiviral proteins, C4 has emerged as a multifunctional pathogenicity determinant that modulates diverse host processes through interactions with numerous host factors. Bhendi yellow vein mosaic virus (BYVMV) (*Begomovirus abelmoschusflavi*) is a monopartite geminivirus associated with a betasatellite that causes yellow vein mosaic disease in okra (14). Similar to several other geminiviral C4 proteins, BYVMV C4 localizes predominantly to the nucleus and plasma membrane and has been implicated in pathogenicity, symptom development, suppression of host antiviral responses, and viral movement (15–17). Considering its functions across distinct subcellular compartments, components of the host nuclear transport machinery are plausible interaction partners that may facilitate C4 functions. A previous affinity purification-mass spectrometry (AP-MS) analysis identified nucleoporin 50a (NUP50a) as a candidate interactor of BYVMV C4 in *Nicotiana benthamiana* (18). Given the established roles of NUP50 proteins in nucleocytoplasmic transport and their emerging nuclear transport-independent functions, we hypothesized that the interaction between C4 and NUP50a may contribute to C4-mediated functions during geminivirus infection. To test this idea, we validated the C4-NbNUP50a interaction using multiple independent approaches and investigated the functional significance of NUP50a for BYVMV infection in *N. benthamiana*. Our findings establish NbNUP50a as a host factor required for efficient BYVMV infection and demonstrate that this protein is targeted by C4 proteins from multiple geminiviruses, suggesting that NUP50a may represent a prevalent host target during geminivirus infection.

## 2. RESULTS

### 2.1 NbNUP50a associates with BYVMV C4 *in planta*

A previous AP-MS analysis of 6x polyhistidine-tagged BYVMV C4 (His-BYVMV C4) identified NUP50a as a candidate host interactor of this viral protein in *N. benthamiana* (18) (Fig. 1A, B). Phylogenetic analysis showed that NUP50 proteins are highly conserved across representative plant species (Fig. S1). Therefore, subsequent analyses focused on NbNUP50a, the orthologue from the experimental host. To validate the interaction, twin-Strep-tagged NbNUP50a (TST-NbNUP50a) and His-BYVMV C4 were transiently expressed individually or together in *N. benthamiana* leaves. Protein extracts were subjected to Ni-NTA pull-down assay. TST-NbNUP50a co-eluted with His-BYVMV C4 only in co-expressed samples, whereas it was detected exclusively in the flow-through fraction when expressed alone, indicating a specific association between the two proteins (Fig. 1C). The interaction was further validated by co-immunoprecipitation from the co-expressed leaf extracts using anti-Strep II antibody. His-BYVMV C4 specifically co-immunoprecipitated with TST-NbNUP50a, whereas no corresponding signal was detected in control samples (Fig. 1D). Together, these results confirm that NbNUP50a associates with BYVMV C4 *in planta*.

**Figure 1.**
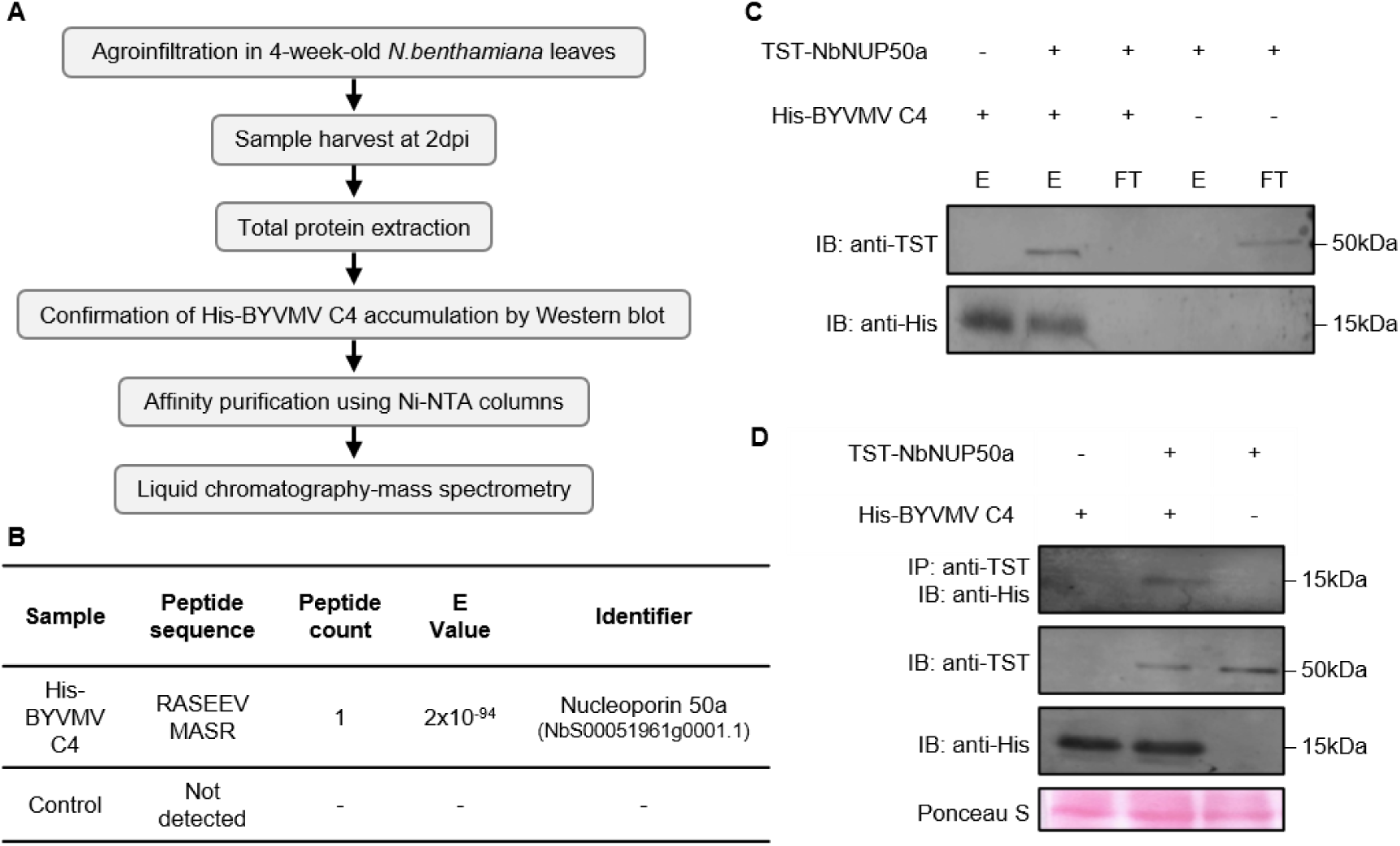
BYVMV C4 associates with NbNUP50a *in planta*. A. Schematic representation of the workflow used for Ni-NTA-based affinity purification coupled with mass spectrometry (AP-MS) in the previous interaction screen (18). B. AP-MS analysis from the previous screening study identified NbNUP50a as a potential interactor of BYVMV C4. C. Ni-NTA-based pull-down assay showing the interaction between 6xHis-tagged BYVMV C4 and twin-Strep-tagged NbNUP50a following transient co-expression in *Nicotiana benthamiana* leaves. E and FT denote the eluted and flow-through fractions, respectively, from the Ni-NTA purification. IP: immunoprecipitation, IB: immunoblot. D. Co-immunoprecipitation of His-BYVMV C4 with twin-Strep-tagged NUP50a following transient co-expression in *N. benthamiana* leaves. Samples expressing His-BYVMV C4 or TST-NUP50a individually were used as negative controls.

### 2.2 BYVMV C4 and NbNUP50a interact directly in the nucleus of the plant cell

A yeast two-hybrid (Y2H) assay was used to test the interaction between BYVMV C4 and NbNUP50a. The results showed that C4 can directly interact with NbNUP50a, as indicated by the growth of yeast cells carrying both constructs on selective media (Fig. 2A). To further examine the subcellular localization of both interacting partners, BYVMV C4-GFP and RFP-NbNUP50a/NbNUP50a-RFP were transiently co-expressed in *N. benthamiana* leaves. Confocal microscopy analysis revealed that both proteins showed strong co-localization in the nucleus (Fig. 2B). In addition, bimolecular fluorescence complementation (BiFC) assays produced fluorescent signals in the nucleus when the fusion proteins containing C4 and NbNUP50a were co-expressed, confirming that the interaction occurs in the nucleus (Fig. 2C).

**Figure 2.**
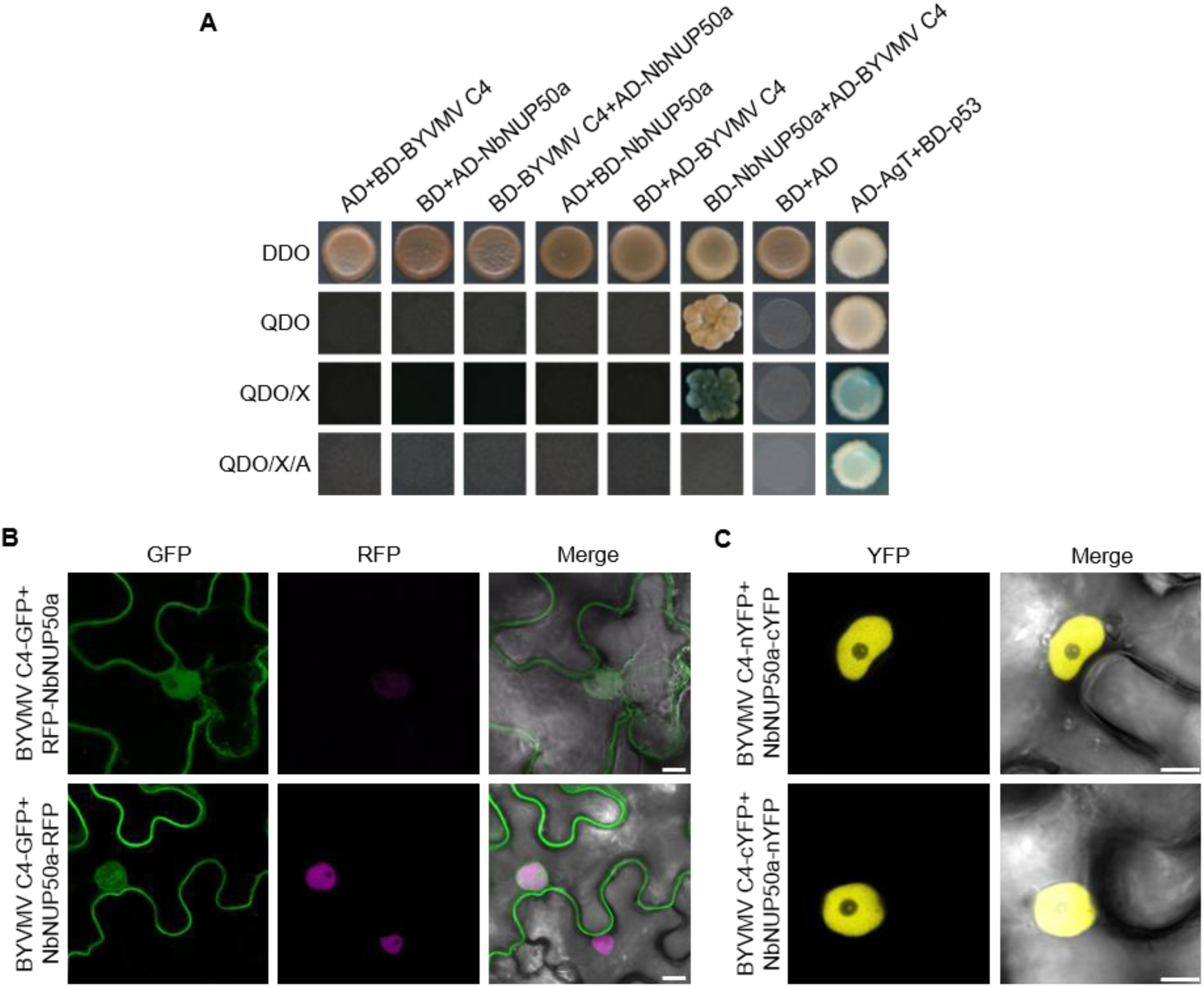
BYVMV C4 interacts directly with NbNUP50a in the nucleus of the plant cell. A. Interaction between BYVMV C4 and NbNUP50a assessed by yeast two-hybrid (Y2H) assay. DDO (double dropout medium): SD/–Leu/–Trp; QDO (quadruple dropout medium): SD/–Ade/–His/–Leu/–Trp; QDO/X: QDO supplemented with X-α-gal; QDO/X/AbA: QDO supplemented with X-α-gal and aureobasidin A. AD-AgT and BD-p53 were used as positive controls for the assay. B. Colocalization analysis of GFP-fused BYVMV C4 with RFP-NbNUP50a or NbNUP50a-RFP. *Agrobacterium tumefaciens* carrying the indicated constructs was infiltrated into *Nicotiana benthamiana* leaves, and fluorescence was imaged at 2 days post-infiltration (dpi). Scale bar: 10 μm. C. Interaction between BYVMV C4 and NUP50a analyzed by bimolecular fluorescence complementation (BiFC) following transient co-expression of indicated constructs in *N. benthamiana* leaves. Images were taken at 2 dpi. Scale bar: 10 μm. YN: N-terminal half of the YFP; YC: C-terminal half of the YFP.

### 2.3 Silencing of *NbNUP50a* reduces BYVMV infection in *N. benthamiana*

To investigate the role of NbNUP50a in BYVMV infection, *NbNUP50a* expression was silenced in *N. benthamiana* using a BYVMV betasatellite-based virus-induced gene silencing (VIGS) vector (hereafter VIGS-EV) (19). A fragment of *NbNup50a* was cloned into the VIGS-EV to generate VIGS-*NbNUP50a*. Three-week-old *N. benthamiana* plants were agroinfiltrated with constructs to express BYVMV, betasatellite, BYVMV+betasatellite, BYVMV+VIGS-EV, BYVMV+VIGS-*NbNUP50a*, and BYVMV+betasatellite+VIGS-*NbNUP50a*. Two independent experimental replicates were performed, and symptom development was recorded in a time-course up to 35 days post-inoculation (dpi) (Fig. 3A; Table 1).

**Figure 3.**
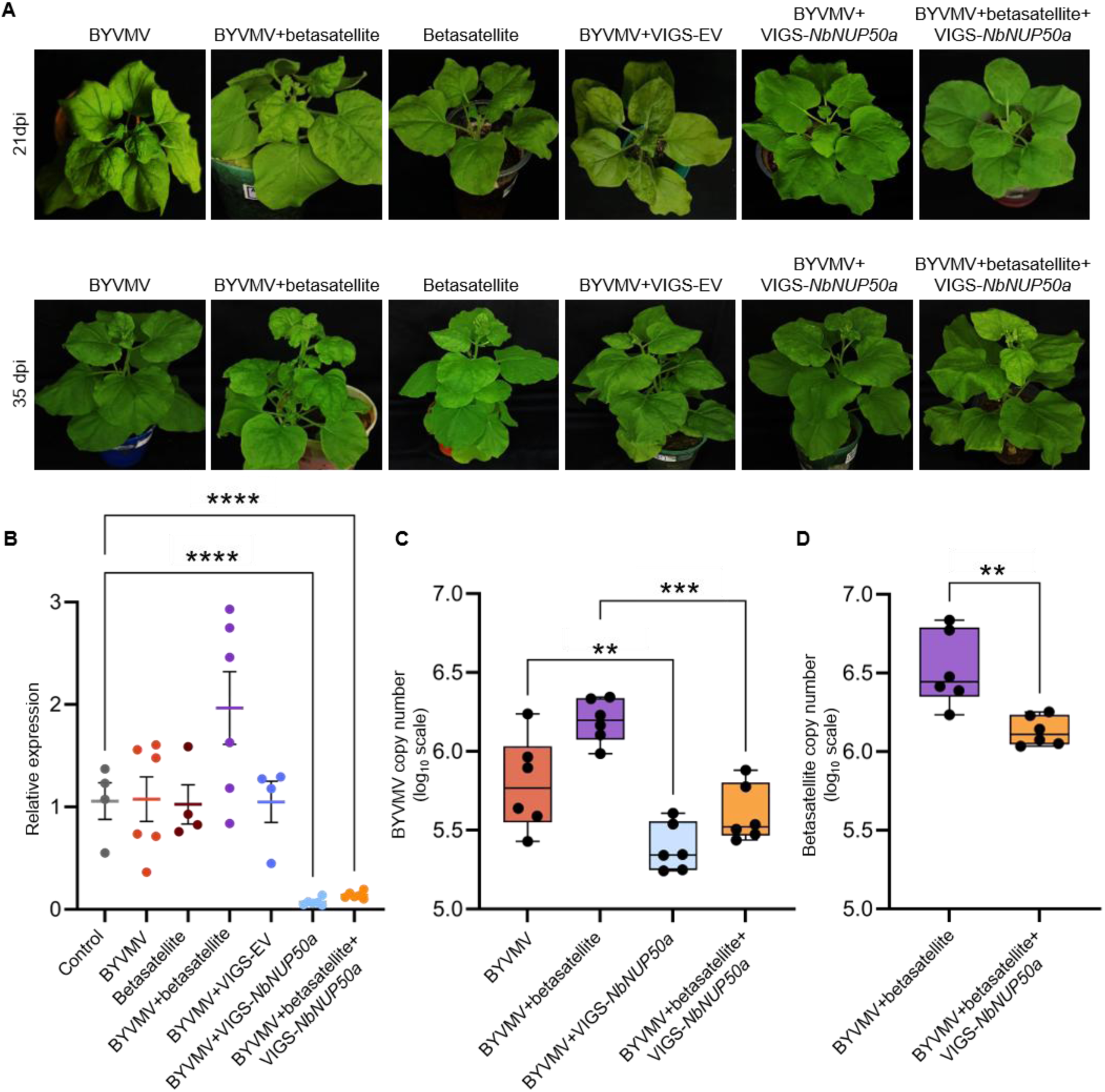
*NbNUP50a* is required for BYVMV infection. A. *NbNUP50a* knockdown in *Nicotiana benthamiana* plants resulted in delayed symptom onset and reduced symptom severity compared with control plants. Three-week-old *N. benthamiana* plants were used for virus-induced gene silencing (VIGS) of *NbNUP50a*. Images in the top and bottom panels were taken at 21 and 35 days post-infiltration (dpi), respectively. B. qRT-PCR analysis confirmed the knockdown of *NbNUP50a* following VIGS. Healthy, uninfiltrated plants were used as controls, and *NbNUP50a* transcript accumulation was quantified in all infiltration combinations relative to the control using the 2^−ΔΔCt^ method. Error bars represent SEM. Each dot represents a biological replicate. Statistical significance was determined from ΔCt values using one-way ANOVA followed by Dunnett’s multiple-comparison test. Asterisks indicate significant differences from the control (P < 0.0001). C. Real-time absolute quantification of BYVMV during infection in control and *NnNUP50a* knockdown plants. BYVMV copy number was calculated using Ct values from a known standard curve and plotted on log₁₀ scale. Error bars represent SEM. Statistical significance was determined using two-way ANOVA with Šídák’s multiple-comparisons test (P < 0.01, P < 0.001). D. Real-time absolute quantification of BYVMV betasatellite during infection in control and *NbNUP50a* knockdown plants. Betasatellite copy number was calculated using Ct values from a known standard curve and plotted on log₁₀ scale. Error bars represent SEM. Statistical significance was analyzed using an unpaired *t*-test with Welch’s correction (P < 0.01).

**Table 1.** Summary of symptoms observed in control and *NbNUP50a*-silenced *N. benthamiana* plants upon BYVMV infection at 35 days post-infection.

| **Combination of viral genome components** | **No. of plants infiltrated** | **Asymptomatic** | **Mild leaf curling and/or stunting** | **Severe leaf curling and/or stunting** |
| --- | --- | --- | --- | --- |
| BYVMV | 20 | 04 | 16 | 0 |
| Betasatellite | 10 | 10 | 0 | 0 |
| BYVMV+betasatellite | 20 | 0 | 03 | 17 |
| BYVMV+VIGS-EV | 10 | 01 | 09 | 0 |
| BYVMV+VIGS-*NbNUP50a* | 24 | 17 | 07 | 0 |
| BYVMV+betasatellite+VIGS-*NbNUP50a* | 24 | 03 | 20 | 01 |

At 21 dpi, plants inoculated with BYVMV+betasatellite developed typical BYVMV symptoms, with symptom onset observed as early as 14 dpi. In contrast, those inoculated with BYVMV and BYVMV+VIGS-EV showed only mild symptoms, while all other combinations remained symptomless. By 35 dpi, BYVMV+betasatellite plants developed severe symptoms, whereas BYVMV and BYVMV+VIGS-EV plants continued to show mild symptoms. All other combinations remained symptomless, except BYVMV+betasatellite+VIGS-*NbNUP50a*, which showed mild leaf curling.

Silencing efficiency was evaluated by reverse transcription quantitative PCR (RT-qPCR) at 21 dpi (Fig. 3B). *NbNUP50a* transcript levels were compared with those of healthy control plants. A significant reduction was observed in BYVMV+VIGS-*NbNUP50a* and BYVMV+betasatellite+VIGS-*NbNUP50a* plants. Expression remained largely unchanged in BYVMV, betasatellite, BYVMV+VIGS-EV, and naive plants, while a slight but non-significant increase was observed in BYVMV+betasatellite plants.

Viral DNA accumulation at 28 dpi was quantified by absolute qPCR using standard curves generated for BYVMV and betasatellite (Fig. 3C, D). A significant reduction in viral copy number was observed in *NbNUP50a*-silenced plants for both genomic components.

### 2.4 *NbNUP50a* silencing does not affect BYVMV C4 localization

Given the role of NUP50a in nucleocytoplasmic transport and the dual localization of BYVMV C4 in nucleus and at plasma membrane, we asked whether NUP50a may contribute to BYVMV infection by regulating the subcellular localization of BYVMV C4. To test this possibility, *NbNUP50a* expression was silenced in *N. benthamiana* using a TRV-based VIGS system, and RT-qPCR confirmed knockdown efficiency at 14 dpi (Fig. S2 A, B). To examine whether *NbNUP50a* silencing affects BYVMV C4 localization, control and *NbNUP50a*-silenced plants were agroinfiltrated with constructs to express BYVMV C4-GFP or free GFP. Subcellular localization of the expressed proteins was analyzed at 2 dpi using confocal laser scanning microscopy. To quantify possible changes in localization, plasma membrane-to-nucleus fluorescence ratios for C4-GFP and cytoplasm-to-nucleus ratios for free GFP were measured using ImageJ. No significant differences in fluorescence distribution or intensity ratios were observed between plants infiltrated with C4-GFP and GFP (Fig. S2C), indicating that *NbNUP50a* knockdown does not alter the subcellular localization of BYVMV C4.

### 2.5 NbNUP50a interacts with C4 proteins from multiple geminiviruses

To evaluate whether the interaction between NUP50a and C4 is conserved among diverse geminiviruses, C4 proteins from okra enation leaf curl virus (OELCuV) (*Begomovirus abelmoschusenation*), TLCYnV, and TYLCV were tested for their ability to interact with NbNUP50a using co-IP, BiFC, colocalization, and Y2H assays. Co-IP assay showed that all three viral C4 proteins associate with NbNUP50a (Fig. 4A). Co-localization analyses using GFP-tagged C4 proteins and NbNUP50a-RFP further demonstrated that the distributions of the tested proteins overlap in the nucleus of the plant cell (Fig. 4B). In addition, BiFC assays confirmed the interaction between all three viral C4 proteins and NbNUP50a, with interaction signals predominantly detected in the nucleus (Fig. 4C). However, a direct interaction with NUP50a in yeast was observed only for TLCYnV C4, as revealed by Y2H analysis (Fig. 4D).

**Figure 4.**
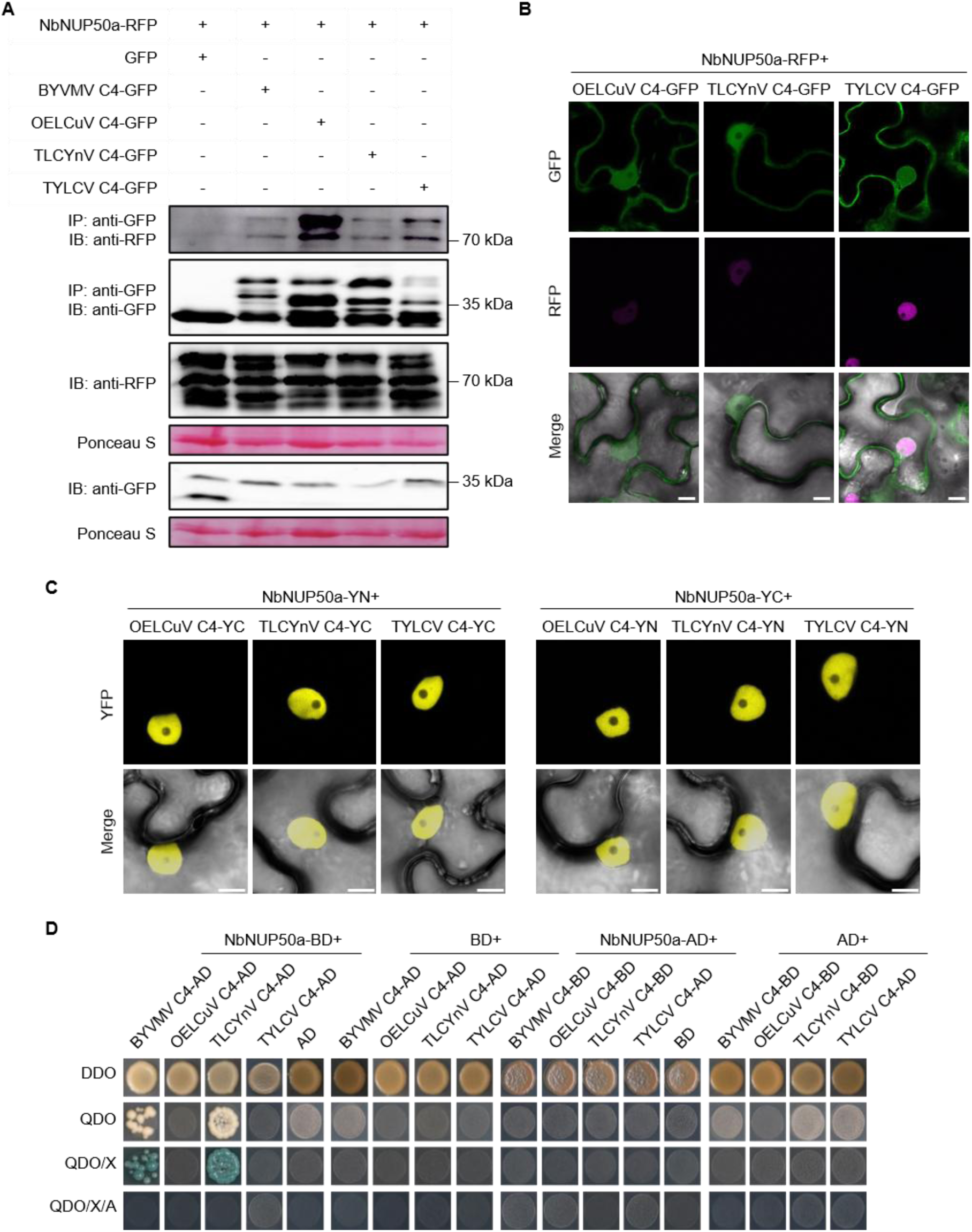
NbNUP50 directly or indirectly interacts with other geminiviral C4 proteins in the nucleus of the plant cell. A. Co-immunoprecipitation of NbNUP50a-RFP with GFP-fused C4 proteins from BYVMV, OELCuV, TLCYnV, or TYLCV following transient co-expression in *Nicotiana benthamiana* leaves. Free GFP was used as a negative control. B. Colocalization analysis of RFP-fused NbNUP50a with OELCuV C4-GFP, TLCYnV C4-GFP, or TYLCV C4-GFP. *Agrobacterium tumefaciens* carrying the indicated constructs was infiltrated into *N. benthamiana* leaves, and fluorescence was imaged at 2 days post-infiltration (dpi). Scale bar: 10 μm. C. Interaction between OELCuV C4, TLCYnV C4, or TYLCV C4 and NbNUP50a analyzed by bimolecular fluorescence complementation (BiFC) following transient co-expression in *N. benthamiana* leaves. Images were taken at 2 dpi. Scale bar: 10 μm. YN: N-terminal half of the YFP; YC: C-terminal half of the YFP. D. Interaction between OELCuV C4, TLCYnV C4, or TYLCV C4 and NbNUP50a assessed by yeast two-hybrid (Y2H) assay. DDO (double dropout medium): SD/–Leu/–Trp; QDO (quadruple dropout medium): SD/–Ade/–His/–Leu/–Trp; QDO/X: QDO supplemented with X-α-gal; QDO/X/AbA: QDO supplemented with X-α-gal and aureobasidin A.

## 3. DISCUSSION

Nucleocytoplasmic trafficking is central to geminivirus infection because viral genomes replicate in the nucleus of infected cells and several viral proteins shuttle between nucleus and cytoplasm to perform distinct functions during the viral cycle. Although several geminiviral proteins interact with host importins and exportins, the role of NPC components in geminivirus infection remains largely unknown. In this study, we identify NbNUP50a as a previously uncharacterized host factor interacting with BYVMV C4 in the nucleus of the plant cell. Functional analyses showed that *NbNUP50a* is required for full BYVMV infection. Interestingly, NbNUP50a interacts with C4 proteins from additional geminiviruses, suggesting that it might be a conserved or convergent target of members of this viral family. Together, these findings establish NUP50a as a nuclear basket nucleoporin that interacts with geminiviral C4 proteins and contributes to successful geminivirus infection (Fig. 5).

**Figure 5.**
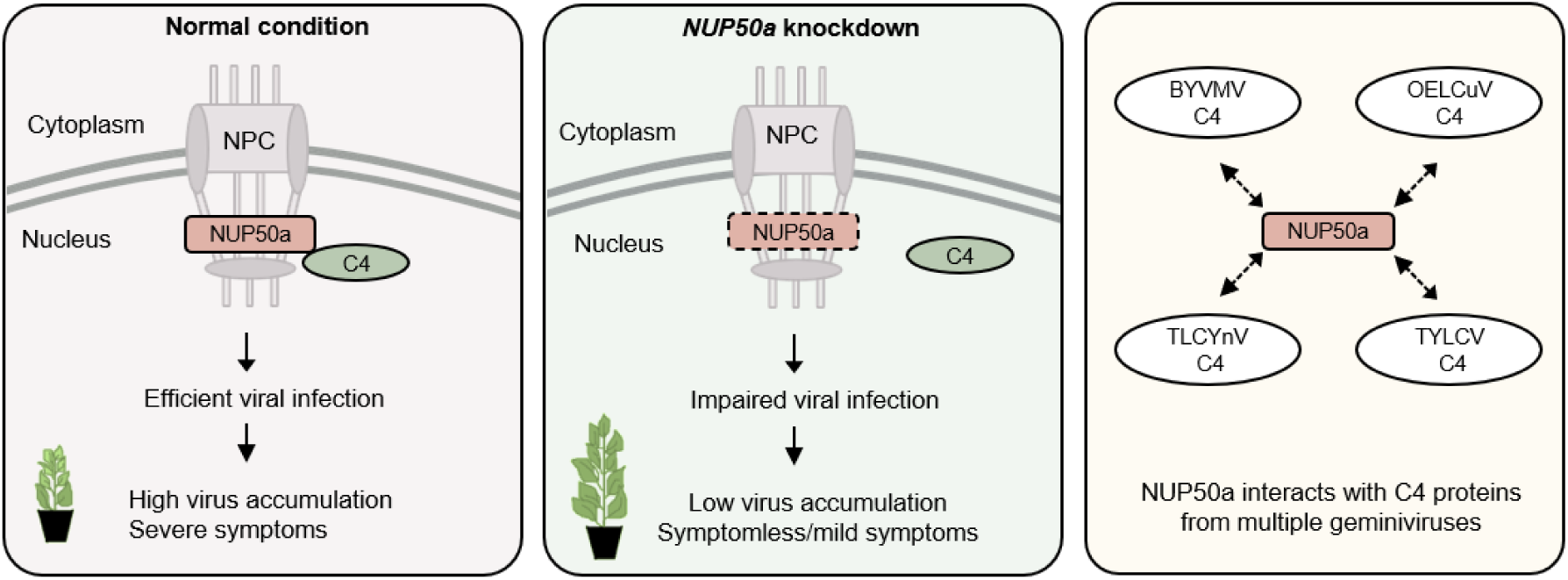
Schematic summary of NUP50a involvement in geminiviral infections. NUP50a, a component of the nuclear pore complex, interacts with geminivirus C4 proteins in the nucleus of the plant cell. Silencing of *NbNUP50a* lead to reduced viral accumulation and no or milder disease symptoms, demonstrating that NUP50a contributes to efficient BYVMV infection. The molecular basis of this contribution remains to be determined but may involve nucleocytoplasmic transport of proteins other than C4 or other NUP50a-dependent cellular processes. The association of NUP50a with C4 proteins from diverse geminiviruses suggests that NUP50a may represent a common host target exploited by diverse geminiviral C4 proteins.

Besides mediating nucleocytoplasmic transport, nucleoporins can also regulate host-pathogen interactions in different ways. In plants, several nucleoporins have been implicated in immune responses. For example, Arabidopsis NUP82 and NUP136 contribute to salicylic acid-dependent immunity against *Pseudomonas syringae* (20), whereas rice NUP98 (APIP12) promotes basal resistance against *Magnaporthe oryzae* (21). In contrast, several nuclear-replicating animal viruses, including members of the *Herpesviridae*, manipulate NPC to gain nuclear entry (22,23) or promote selective import of viral proteins (24). Together, these studies show that nucleoporins can either promote host defense or be exploited by pathogens, depending on the biological context. However, whether they contribute to infection by plant DNA viruses has remained unknown. Our findings represent a first step towards bridging this gap by demonstrating that a nuclear basket nucleoporin can function as a susceptibility factor during geminivirus infection.

NUP50a is a highly dynamic FG-repeat nucleoporin that interacts with importin α, importin β, and Ran GTPases during cargo transport (7). Unlike scaffold nucleoporins, NUP50a spends only a short time associated with the NPC and is highly mobile within the nucleoplasm, allowing it to function at the interface between the NPC and the nuclear transport machinery (25,26). BYVMV C4 exhibits a dual localization to the nucleus and the plasma membrane, indicating that it may undergo dynamic intracellular trafficking. Since nuclear accumulation requires transport across the nuclear envelope through the NPC, we hypothesized that NUP50a might regulate the subcellular localization of C4 by facilitating its nucleocytoplasmic transport. However, although silencing of *NbNUP50a* significantly reduced viral accumulation, it did not alter the subcellular localization of BYVMV C4. These results suggest that NUP50a does not simply determine C4 localization; instead, it may facilitate other transport-related events during infection. Further research will be required to uncover the exact molecular mechanism underlying the positive contribution of NUP50a, and of its interaction with C4, to the viral infection.

In bipartite geminiviruses, nucleocytoplasmic trafficking of viral DNA is mediated by the nuclear shuttle protein (NSP) (27). In contrast, monopartite geminiviruses lack NSP. Instead, the CP has been implicated in intracellular trafficking of viral genomes, with nuclear export being facilitated by proteins such as V2 in several monopartite geminiviruses (28). Our observation that NbNUP50a interacts with BYVMV C4, yet does not alter its steady-state localization, raises the possibility that NUP50a facilitates the assembly or trafficking of viral DNA-protein complexes rather than directing C4 localization itself. Alternatively, given its established role in nucleocytoplasmic transport, NUP50a may regulate the nuclear transport of specific host factors that are required for efficient viral infection. Impaired trafficking of such host cargoes could also account for the marked reduction in viral accumulation observed in *NbNUP50a*-silenced plants.

NUP50a may also promote infection through transport-independent functions. Increasing evidence indicates that several nucleoporins are involved in transcriptional regulation and RNA processing. Recent proximity-labeling studies identified AtNUP50a in complexes enriched with spliceosomal proteins and transcriptional regulators (29). In addition, NUP50 homologues have been linked to chromatin organization, maintenance of chromatin boundaries, and regulation of actively transcribed genes (30,31). Since geminiviruses extensively reprogram host transcription during infection (32), C4-mediated recruitment of NUP50a may influence transcriptional or post-transcriptional processes that favour infection.

NUP50a also associated with C4 proteins from three additional geminiviruses, suggesting that targeting of NUP50a may represent a prevalent strategy among geminiviruses. Although only BYVMV C4 and TLCYnV C4 showed a detectable direct interaction in yeast, all four C4 proteins associated with NUP50a *in planta* by co-IP and BiFC. These findings suggest that the molecular basis of the NUP50a-C4 interaction may differ among geminiviruses, with some C4 proteins binding NUP50a directly and others associating through additional host factors or protein complexes. Such differences may reflect independent evolutionary adaptations that converge on the same host target.

Although our findings establish that NUP50a promotes geminivirus infection, the precise role of the C4-NUP50a interaction remains unclear. Future studies should determine whether NUP50a contributes to viral replication, movement, or modulation of host antiviral responses. Mapping the interaction domains and identifying proteins associated with the C4-NUP50a complex will help further define its molecular function.

Overall, this study provides evidence linking a nuclear basket nucleoporin to geminivirus pathogenesis and identifies NUP50a as a common host target of C4 proteins from multiple geminiviruses. These findings provide a foundation for understanding how nuclear pore-associated processes contribute to plant DNA virus infection.

## 4. MATERIALS AND METHODS

### Plasmids and cloning

*Escherichia coli* strains DH5α and Top10 were used for general cloning procedures, while DB3.1 was used to amplify the Gateway-compatible empty vectors. The constructs generated in this study and their cloning strategy are listed in Supplementary Table 1.

ORF coding for *NbNUP50a* (NbS00051961g0001.1) was amplified using the corresponding primers (Supplementary Table 1) and cloned into pDONR/Zeo entry vector through BP reactions as part of Gateway cloning. LR reactions were performed to recombine them into pGWB554, pGWB555 (33), pGTQL1211YC, and pGTQL1211YN (34) to generate NbNUP50a-RFP, RFP-NbNUP50a, NbNUP50-YC, and NbNUP50-YN. His C4, TST-NbNUP50a, AD-/BD-NbNUP50a, VIGS-*NbNUP50a,* and TRV2-*NbNUP50a* were generated through cloning the corresponding ORFs in pGA643, pGADT7, pGBKT7, VIGS-EV (19), and TRV2 by restriction-ligation or in-fusion cloning. The details of primers used are listed in Supplementary Table 1.

pGWB505-/pGTQL1211YC-/pGTQL1211YN/pGADT7-/pGBKT7-BYVMV C4, pGWB505-/pGTQL1211YC-/pGTQL1211YN-/pGADT7-/pGBKT7-OELCuV C4, pGWB505-/pGTQL1211YC-/pGTQL1211YN/pGADT7-/pGBKT7-TLCYnV C4, and the binary vectors to express TYLCV C4-GFP, TYLCV C4-nYFP/cYFP, and AD-/BD-TYLCV C4 were previously described (17,35,36).

### Plant growth conditions

*N. benthamiana* plants were grown in controlled growth rooms under a 16 h light/8 h dark cycle (long-day conditions) at 25°C.

### Agroinfiltration in *N. benthamiana*

Corresponding constructs were mobilized into *Agrobacterium tumefaciens* strain EHA105 or GV3101 by triparental mating or heat-shock transformation as described in (37), with minor modifications. Transformants or transconjugants were cultured in Luria-Bertani liquid medium overnight at 28°C and were pelleted down by centrifugation at 4000 g at room temperature for 10 min, followed by resuspension in infiltration buffer (10 mM MgCl2, 10mM MES [pH 5.6], and 150 μM acetosyringone). The final OD600 was adjusted to 0.1-0.5 and maintained in the dark for 2-4 hours before infiltration. Equal volumes of each construct were mixed in the case of co-infiltrations. The bacterial suspensions were infiltrated into the abaxial side of fully expanded young leaves in 3-4-week-old *N. benthamiana* plants using a 1 ml needleless syringe.

### Virus-induced gene silencing (VIGS) in *N. benthamiana* plants

VIGS-mediated silencing was carried out using either a BYVMV betasatellite-derived vector (VIGS-EV) or a tobacco rattle virus (TRV)-based system. A 425 bp gene fragment for silencing *NbNUP50a* was selected using the Sol Genomics Network VIGS tool (38) to minimize potential off-target effects. For the betasatellite-based system, *A. tumefaciens* cultures carrying VIGS-EV-derived constructs were co-infiltrated with the APTR helper construct. For TRV-mediated silencing, *A. tumefaciens* cultures harbouring TRV2-derived constructs were mixed with TRV1 cultures. Following agroinfiltration, plants were maintained under controlled growth conditions, and silencing efficiency was evaluated by RT-qPCR at 14–21 days post-infiltration.

### Protein extraction, pull-down, and co-immunoprecipitation assay

Approximately 1 g of leaf discs from infiltrated *N. benthamiana* plants was harvested at 2 dpi. Total protein was extracted using a buffer including 50 mM Tris-HCl pH 7.5, 150 mM NaCl, 10% glycerol, 10 mM DTT, 10 mM EDTA, 1% (v/v) protease inhibitor cocktail (Sigma), and 1% (v/v) IGEPAL CA-630 (Sigma).

For the affinity purification-based pull-down assay, Ni-NTA agarose resin columns were used following the manufacturer’s protocol (HiMedia). The columns, post-incubation with protein lysates, were washed with 25-40mM imidazole, and the final protein complex was competitively eluted with 250-400mM imidazole. Co-immunoprecipitation was performed as described previously (39), with minor modifications. For immunoprecipitation with anti-Strep II, the antibody was conjugated with Dynabeads Protein A (Thermo Scientific, 10001D) before incubation with protein lysates. The primary and secondary antibodies used are the following: mouse anti-Strep II (Abcam, GT661; 1:1000 dilution), mouse anti-His (Thermo Scientific, MA1-21315; 1:2000 dilution), mouse anti-RFP (ChromoTek, 6G6; 1:5000 dilution); goat anti-GFP (SICGEN, AB0020-500; 1:5000 dilution); anti-mouse HRP (Invitrogen, 62-6520; 1:15000 dilution), anti-mouse peroxidase (Sigma-Aldrich, A2554; 1:15000 dilution); anti-goat peroxidase (Sigma-Aldrich, A8919; 1:20000 dilution).

### Confocal microscopy

Confocal imaging was performed on a Zeiss LSM880 Upright confocal laser scanning microscope using the pre-set parameters for GFP with excitation (Ex) 488 nm and emission (Em) 500-550 nm, and for RFP with Ex 561 nm and Em 580-630 nm. Leaf discs of *N. benthamiana* plants transiently expressing the fluorescence-tagged proteins were imaged at 2 dpi.

### Bimolecular fluorescence complementation assay

Leaf discs from *N. benthamiana* plants agroinfiltrated with constructs to transiently express the corresponding proteins were visualized at 2 dpi. Imaging was performed on a Zeiss LSM880 Upright laser scanning confocal microscope using the preset parameters for YFP with Ex 514 nm and Em 525–575 nm.

### Yeast-two-hybrid assay

All yeast constructs were introduced into the *Saccharomyces cerevisiae* Y2HGold strain (Clontech) using the Frozen-EZ Yeast Transformation II Kit (Zymo) according to the manufacturer’s protocol. Subsequent selection of transformants and assessment of protein-protein interactions on selective media were carried out as described previously (35). Briefly, colonies that grew on minimal synthetic defined (SD) medium lacking leucine and tryptophan (double dropout, DDO) were collected and resuspended in 20 μL of liquid DDO medium. Aliquots of 3-4 μL from each suspension were then spotted onto SD plates lacking leucine, tryptophan, histidine, and adenine (quadruple dropout, QDO), QDO supplemented with X-α-gal (QDO/X), and QDO/X containing aureobasidin A (QDO/X/AbA). The plates were incubated at 28 °C in the dark, and colony growth and color development were documented by photography after 5-6 days.

### RNA extraction and RT-qPCR

RNA was extracted using RNA extraction reagent RNAiso Plus (Takara) or by the citrate-citric acid method (40), followed by DNase I (Thermo Scientific) treatment to remove genomic DNA. 500 ng of RNA was converted to cDNA using RevertAid First Strand cDNA Synthesis kit (Thermo Scientific) or PrimeScript RT MasterMix (Takara), following the manufacturer’s instructions. qPCR reactions were run with PowerTrack SYBR Green Mastermix (Thermo Scientific), diluted cDNA template, and corresponding primers (Supplementary Table 2), in a QuantStudio 1 (Thermo Scientific) with the following program: 2 min at 95°C, and 40 cycles consisting of 15 s at 95°C, 1 min at 60°C. *NbActin* was used as a normalizer gene. Comparative analyses of transcripts were performed by applying the 2^−ΔΔCt^ method.

### Viral DNA extraction and quantification

BYVMV A DNA (AF241479) and betasatellite (AJ308425) were used in the infection studies (14). Viral DNA was extracted from infected leaf tissue using the CTAB-alkaline lysis method as previously described (41). Briefly, approximately 1 g of leaf tissue was ground in liquid nitrogen, mixed with 2x CTAB buffer, and incubated at 60 °C for 10 min. DNA was purified by chloroform:isoamyl alcohol extraction and ethanol precipitation, followed by selective enrichment of circular viral DNA using SDS, NaOH, and sodium acetate treatments. The final DNA pellet was resuspended in sterile molecular-grade water. qPCR reactions containing equal concentrations of viral DNA as template, PowerUp SYBR Green Master Mix, and gene-specific primers (Supplementary Table 2) were run on a QuantStudio 1 with the following program: 2 min at 95°C, and 40 cycles consisting of 15 s at 95°C, 1 min at 60°C. Standard curves were generated using ten-fold serial dilutions of plasmids containing viral genome inserts, and viral DNA copy numbers were calculated from the corresponding Ct values.

### Fluorescence intensity analysis

Fluorescence intensity ratios were calculated using ImageJ, following a previously described method (42), to evaluate changes in the subcellular distribution of GFP and BYVMV C4-GFP in control and *NbNUP50a*-silenced plants. The nucleus-to-cytoplasmic fluorescence intensity ratio was used for GFP, whereas the nucleus-to-plasma membrane fluorescence intensity ratio was used for C4-GFP to quantify their relative abundance in different cellular compartments.

## ACKNOWLEDGEMENTS

The authors thank Huang Tan, Laura Medina-Puche, Shaojun Pan, and Hua Wei for critical reading of the manuscript and help with figure design, and Bettina Stadelhofer and the central facilities at the ZMBP, especially the Plant Cultivation and the Microscopy facilities, for excellent technical support. The authors also acknowledge Jeyalakshmi Karanthamalai for initial discussions and Deepanjan Banerjee for assistance with preliminary experiments. AC acknowledges support from the DAAD bi-nationally supervised doctoral programme and DST-INSPIRE Fellowship. Work in the RLD lab was partially funded by the DFG (TRR 356/1 2023 – 491090170) and the European Research Council (GemOmics; 101044142). Work in the GP lab was partially supported by DBT (Ref. No. BT/PR23641/BPA/118/309/2017) and SERB (Ref. No. EEQ/2022/000909).

## SUPPLEMENTARY FIGURES

**Figure S1.**
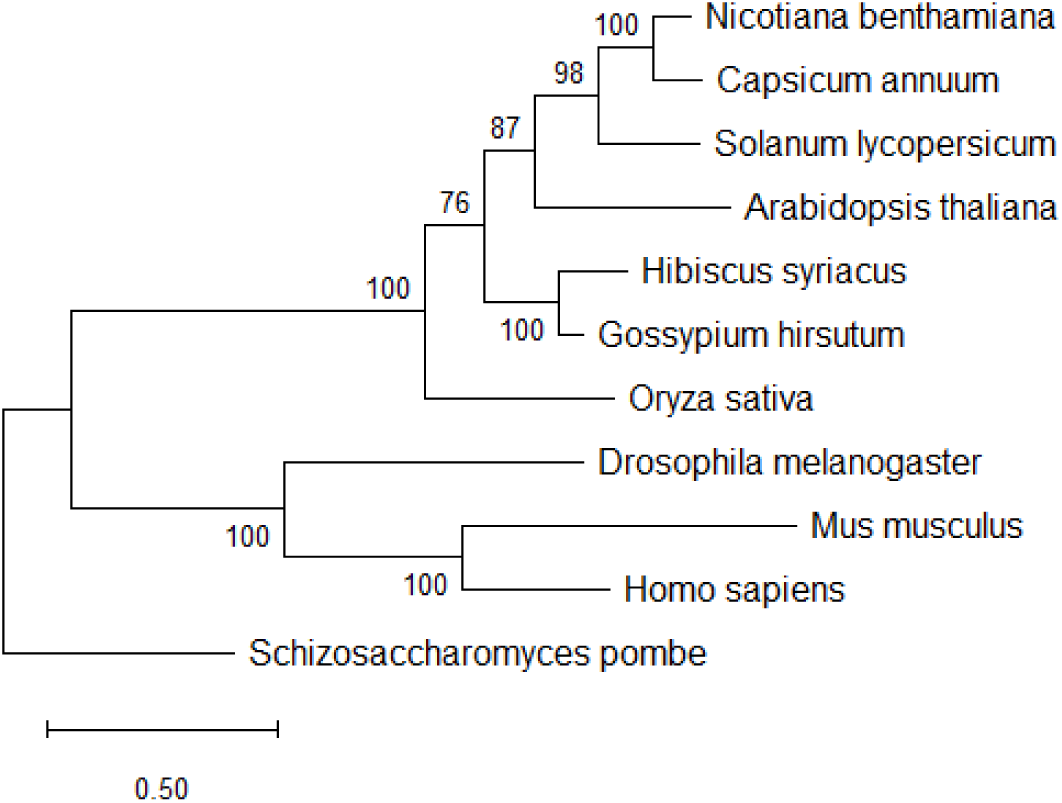
Phylogeny of NUP50a. Maximum likelihood phylogenetic tree of NUP50a homologs constructed using MAFFT alignment and MEGA software with 1000 bootstrap replicates. Bootstrap values are indicated at branch nodes. *Schizosaccharomyces pombe* was used as the outgroup.

**Figure S2.**
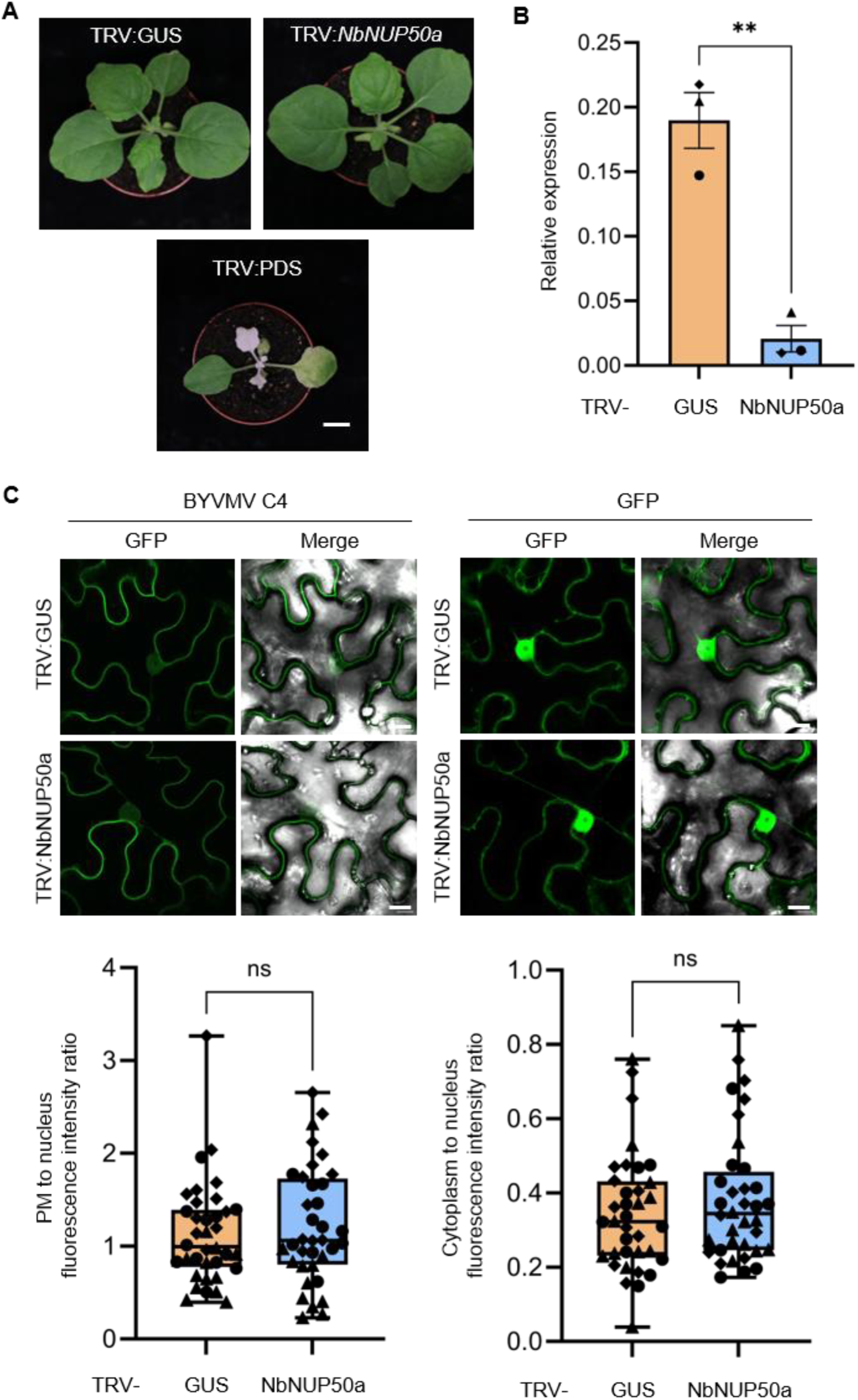
Silencing of *NbNUP50a* does not affect the subcellular localization of BYVMV C4. A. *Nicotiana benthamiana* plants were silenced for *NbNUP50a* expression through TRV-based VIGS. Scale bar: 3 cm. B. RT-qPCR analysis confirming *NbNUP50a* silencing. Bars represent the mean ± SEM, and each symbol represents a biological replicate. *NbActin* was used as the reference gene for normalization. C. Control and *NbNUP50a*-silenced *N. benthamiana* plants were agroinfiltrated with BYVMV C4-GFP or free GFP constructs for transient expression. Confocal images were captured at 2 dpi. Scale bar: 10 μm. Fluorescence intensity ratios were quantified using ImageJ as plasma membrane-to-nucleus ratios for BYVMV C4-GFP and cytoplasm-to-nucleus ratios for free GFP. Data were obtained from three independent biological replicates. Each point represents an individual cell, and identical symbols denote cells from the same plant. No significant differences were observed between control and silenced plants according to Welch’s t-test.

